# Transporter genes in biosynthetic gene clusters predict metabolite characteristics and siderophore activity

**DOI:** 10.1101/2020.06.24.170084

**Authors:** Alexander Crits-Christoph, Nicholas Bhattacharya, Matthew R. Olm, Yun S. Song, Jillian F. Banfield

## Abstract

Biosynthetic gene clusters (BGCs) are operonic sets of microbial genes that synthesize specialized metabolites with diverse functions, including siderophores and antibiotics, which often require export to the extracellular environment. For this reason, genes for transport across cellular membranes are essential for the production of specialized metabolites, and are often genomically co-localized with BGCs. Here we conducted a comprehensive computational analysis of transporters associated with characterized BGCs. In addition to known exporters, in BGCs we found many importer-specific transmembrane domains that co-occur with substrate binding proteins possibly for uptake of siderophores or metabolic precursors. Machine learning models using transporter gene frequencies were predictive of known siderophore activity, molecular weights, and a measure of lipophilicity (log *P*) for corresponding BGC-synthesized metabolites. Transporter genes associated with BGCs were often equally or more predictive of metabolite features than biosynthetic genes. Given the importance of siderophores as pathogenicity factors, we used transporters specific for siderophore BGCs to identify both known and uncharacterized siderophore-like BGCs in genomes from metagenomes from the infant and adult gut microbiome. We find that 23% of microbial genomes from the infant gut have siderophore-like BGCs, but only 3% of those assembled from adult gut microbiomes do. While siderophore-like BGCs from the infant gut are predominantly associated with *Enterobactericaee* and *Staphylococcus*, siderophore-like BGCs can be identified from taxa in the adult gut microbiome that have rarely been recognized for siderophore production. Taken together, these results show that consideration of BGC-associated transporter genes can inform predictions of specialized metabolite structure and function.

## Introduction

Microbes produce specialized metabolites with diverse functions, including siderophores, ionophores, antibiotics, antifungals, and signalling molecules (Osbourn 2010). Specialized metabolites therefore often underlie both cooperative and competitive interactions between microbes, and microbial interactions with the physiochemical environment (Davies 2013; Sharon et al. 2014; Tyc et al. 2017). The vast majority of specialized metabolites in bacteria are produced by biosynthetic gene clusters (BGCs), which are sets of genomically colocalized genes. Colocalization of genes into BGCs is thought to occur because of selection for co-inheritance and co-regulation (Fischbach, Walsh, and Clardy 2008). While thousands of microbial natural products have been characterized, genomic BGC predictions made using programs such as antiSMASH (Blin, Shaw, et al. 2019) and ClusterFinder (Cimermancic et al. 2014) suggest that characterized molecules represent just a small fraction of all existing microbial natural products (Medema and Fischbach 2015; Kim et al. 2017). Many of these unknown metabolites may be highly novel due to enzymatic and combinatorial diversity of genes in BGCs (Jenke-Kodama, Börner, and Dittmann 2006; Chevrette et al. 2019).

Because of the sheer number of sequenced but otherwise uncharacterized BGCs and the time and costs required for chemical characterization, there is a pressing need for predictions of BGC metabolite structures or functions to enable prioritization of targets for laboratory study (Tran et al. 2019). Prediction of metabolite structure or function for a novel BGC from gene content alone is challenging. For many biosynthetic nonribosomal peptide synthetases (NRPS) and polyketide synthases (PKS), there is a ‘co-linear’ assembly-line regulation in which the order of genes relates to the order of enzymatic modifications on the metabolite during synthesis (Fischbach and Walsh 2006). Using this co-linearity rule can help predict some degree of structural detail in NRPSs and PKSs, as is done by by antiSMASH and PRISM (Skinnider et al. 2017), but there are many known exceptions to this rule (Wenzel and Müller 2005), and the accuracies of these software predictions have not been formally assessed using a large training dataset.

Prediction of BGC metabolite function generally relies on contextual genes associated with BGCs. The observation that genes conferring resistance to the produced metabolite are also colocalized with the BGC motivates investigation of putative resistance genes (self-resistance gene mining (Yan, Liu, and Tang 2020)) for functional prediction. For siderophore activity prediction, AntiSMASH assigns a functional ‘siderophore’ label for BGCs that contain the IucA / IucC gene family, but this gene is only specific for siderophores with biosynthetic pathways similar to aerobactin (Hider and Kong 2010). More recently, Hannigan et al. (2019) trained neural networks to both identify BGCs in genomes and classify BGCs by known metabolite functions. These networks used Protein Families from the PFAM database (“PFAM Database,” n.d.) (PFAMS) found in each BGC as features. They predicted activity labels of antibacterial, antifungal, cytotoxic, and inhibitor, with precisions of 36%, 47%, 61%, and 69% on each class respectively.

Many specialized metabolites perform their ecological roles extracellularly, and thus require transport across cellular membranes. Transporter genes often colocalize in BGCs and have been shown to be compound specific and necessary for export of the product in many cases (Severi and Thomas 2019; Martín, Casqueiro, and Liras 2005; Méndez and Salas 2001). Therefore, transporters may also inform predictions of BGC metabolite structure and function. The distribution of transporters associated with biosynthetic gene clusters has so far been assessed only in characterized BGCs with experimental validation, a small fraction of the total number BGCs sequenced. At least 40 BGC-associated exporters have been characterized, mostly in the *Actinomycetes*, with varying degrees of experimental validation (Severi and Thomas 2019).

Transporters associated with BGCs are commonly either ATP-dependent active transporters or ion-gradient dependent transporters (Martín, Casqueiro, and Liras 2005). ATP-dependent transporters include the ATP-binding cassette (ABC) superfamily of both importers and exporters (Rees, Johnson, and Lewinson 2009), and the MacB tripartite efflux pump (Greene et al. 2018b). Examples of characterized structures of each transporter class and their substrates are shown in **Fig 1a**. In brief, Type I ABC importers are characterized by the *BPD_transp_1* transmembrane (TM) protein family, and include MalFGK and MetNI for malate and methionine import in *E. coli* (Beek et al. 2014). Type II ABC importers are characterized by the *FecCD* TM protein family, and examples include BtuCD and HmuUV (Beek et al. 2014), and the FecBCDE system for Iron(III) dicitrate import in *Escherichia coli* (Staudenmaier et al. 1989). Both types of ABC importers often associate with Substrate Binding Proteins (SBPs), small membrane or periplasmic proteins for substrate uptake (Beek et al. 2014; Berntsson et al. 2010). Periplasmic binding proteins, Type II ABC importers, and TonB-dependent receptors are also known to play key roles in siderophore uptake in multiple bacterial species (Ellermann and Arthur 2017).

**Figure 1:**
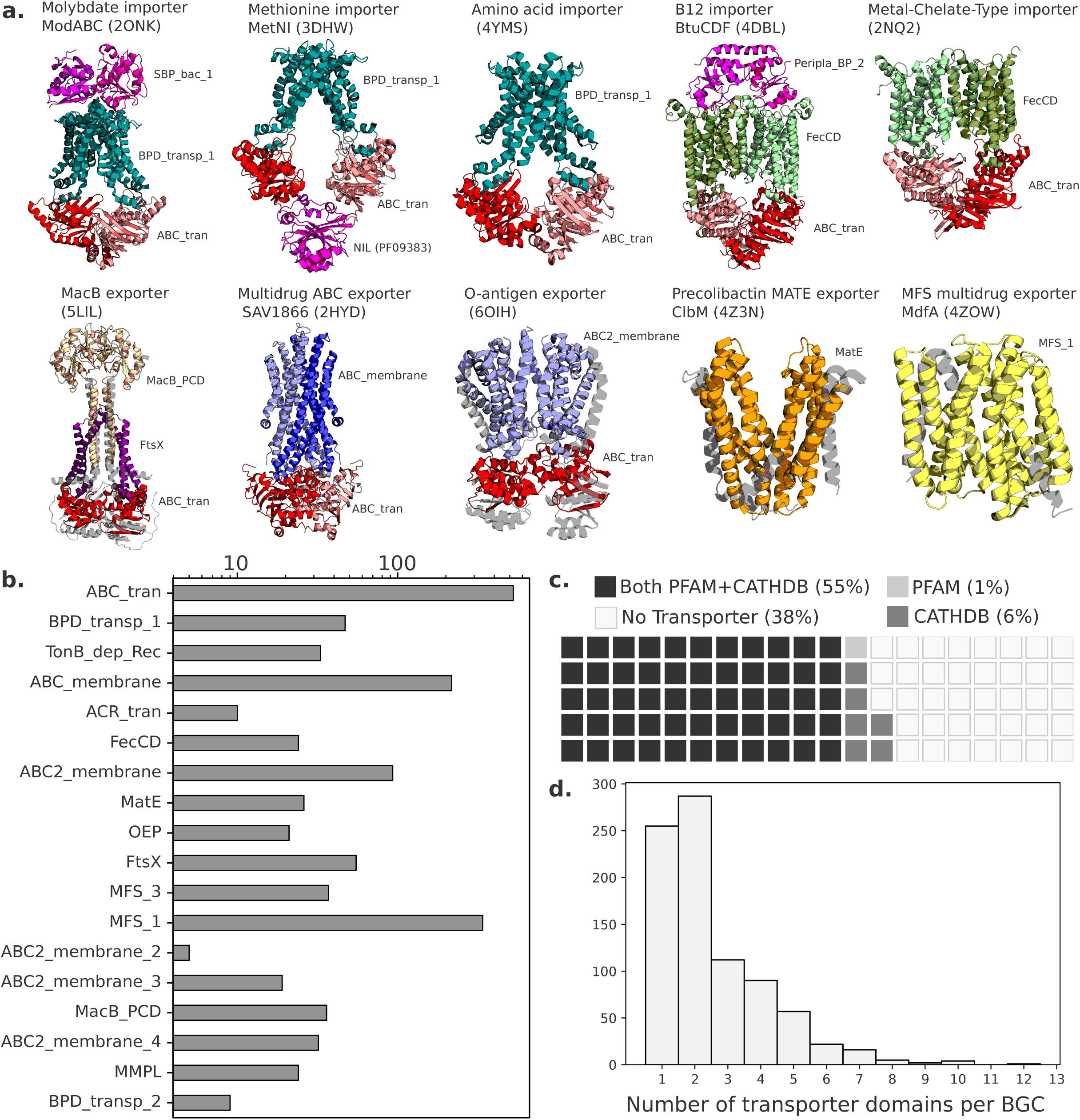
Distributions of transporter classes in biosynthetic gene clusters. (a) Structures of characterized examples of major transporter classes often found in olored and labeled by PFAM domains. (b) The frequencies of common PFAM rter domains across the bacterial BGCs in the MIBiG database. (c) The Percentages of bacterial BGCs in MiBiG that do and do not contain transporter domains. Each square represents 1% of BGCs. (d) The counts of transporter domains per each bacterial BGC that contains at least 1 transporter gene across MiBiG.

Meanwhile, examples of ABC exporters include McjD for lasso peptide microcin J25 export (Romano et al. 2018) and the *Staphylococcus aureus* multidrug exporter Sav1866 (Dawson and Locher 2007), composed of the *ABC_mem* TM protein family, while the O-antigen polysaccharide exporter is composed of the *ABC2_membrane* TM protein family (Bi et al. 2018). Export of Nystatin, Doxorubicin, and Mccj25 was found to be dependent upon ATP-dependent transporters (Severi and Thomas 2019). The vast majority of ribosomally synthesized and post-translationally modified peptides (RiPPs) and a number of antibiotics from *Actinomycetes* also rely on characterized ATP-dependent ABC transporters (Gebhard 2012) (Méndez and Salas 2001).

Ion-gradient dependent transporters (also known as secondary active transport systems) do not require ATP and facilitate transport of small molecules in response to chemiosmotic gradients (Quistgaard et al. 2016). Those found in BGCs are often examples of the major facilitator superfamily (MFS) and occasionally, the resistance nodulation division (RND) family or the multidrug and toxic compound extrusion (MATE) family. Examples of characterized secondary active transporters for antibiotics include an RND transporter for pyoluteorin, and MFS transporters specific for mitomycin C, virginiamycin S, and landomycin (Severi and Thomas 2019).

Thousands of BGCs with chemically characterized metabolites open up the possibility for a broad genomic and computational analysis of phylogenetically and functionally diverse BGCs. Here, we used a curated version of the Minimum Information about a Biosynthetic Gene cluster database (MIBiG 2.0) of BGCs (Kautsar et al. 2020) and selected transporter-specific protein domain hidden markov models (HMMs) to perform a wide genomic assessment of the distribution of transporters in BGCs. We found clear correlations between transporter domains and corresponding metabolite features, especially siderophore activity, that indicate underlying logical structure to transporter associations and can inform functional and structural prediction of specialized metabolites from genomics alone.

## Results

### Genome mining of transporters associated with biosynthetic gene clusters

Using two compiled sets of transporter-specific HMMs (PFAM and CATHDB), we cataloged all classes of transporters across the MIBiG 2.0 database of characterized and experimentally validated biosynthetic gene clusters. We found that 56% of the bacterial BGCs in MIBiG contained at least one PFAM transporter hit and an additional 6% contained a CATHDB transporter hit without a PFAM domain (**Fig 1c**). These percentages increased among BCGs that produce antibiotics (71%) and siderophores (78%), indicating that BGCs with these activities are more likely to contain at least one transporter. BGCs with transporters contained 2.5 transporter-associated domains across transport-annotated genes on average (**Fig 1d**), which is expected as many ATP-dependent transporter systems have at least 2 domain complexes. However, some BGCs contained considerably more transporter ORFs and domains, indicating that sometimes multiple transport systems can be associated with one BGC (**Supplementary Tables S2-S3**). The number of transporters in a BGC had no association with the number of metabolite structures reported for that BGC. The ATP-binding ABC transporter (*ABC_tran*) domain and the Major Facilitator Superfamily 1 (*MFS_1*) domain were the two most common transporter domains found in BGCs (**Fig 1b**). A variety of proteins had nucleotide-binding domains along with several different transmembrane domains – *ABC_membrane* and *ABC2_membrane* domains were most common but *ABC2_membrane_2*, -*_3*, and -*_4* domains were also represented.

Examining domains specific for export, the *ABC_membrane* domain is often characteristic of exporters (e.g., Sav1866 (Velamakanni et al. 2008)), but recently has been reported in the genes for siderophore uptake (YbtPQ) in *Yersinia (Wang, Hu, and Zheng 2020)*, and is therefore not necessarily indicative of export or import alone. The second most common transmembrane domain, *ABC2_membrane*, has been observed in the O-antigen polysaccharide exporter (Bi et al. 2018). The *MacB-FtsX* tripartite efflux pump was found in 60 BGCs, while the RND (*ACR_tran*) efflux pump was less common, and found in only 10 BGCs. Other known efflux systems, such as SMR, MatE, and the MFS families 2 through 5 were comparatively rare across BGCs.

We next calculated co-occurrence correlations between all transporter protein families across BGCs and observed a strong negative correlation between *MFS* transporters and ATP-dependent transporters relying on the nucleotide binding domain, and a weaker negative correlation of *MatE* domains from the ATP-dependent NBD (**Fig 2a**). This points towards a dichotomous choice between ATP-dependent and ATP-independent transport associated with a BGC.

**Figure 2:**
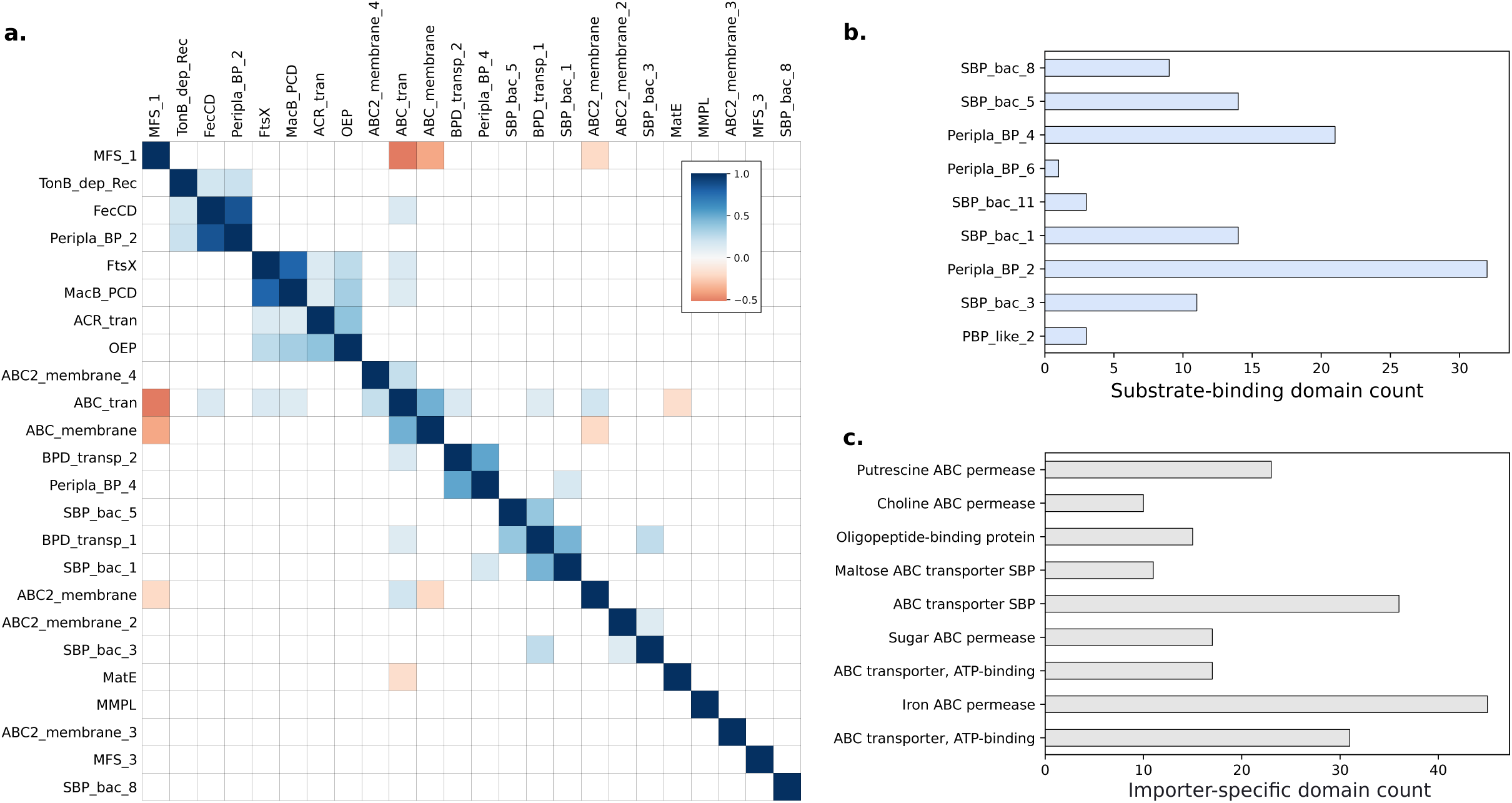
Presence of importer-specific domains and co-occurrence between transporters across BGCs. (a) Spearman correlations between commonly occurring PFAM transporter domains across MiBiG BGCs - only correlations with p < 0.001 are shown. (b) Counts of PFAM transporter substrate binding domain families, and (c) and counts of importer-specific CATHDB domains across MiBiG BGCs. The CATHDB functional family “Iron ABC permease” is essentially synonymous with the FecCD PFAM.

Multiple lines of annotation evidence indicated that many of the transporter genes associated with BGCs were likely to be importers. Importers can be involved in the uptake or re-uptake of molecules like siderophores, and may also play roles in importing precursor metabolites for a BGC. The membrane domains specific to Type I importers (*BPD_transp_1*) and Type II importers (*FecCD*) were the most often observed ATP-dependent transmembrane domains besides *ABC_membrane* and *ABC2_membrane* (**Fig 1b**). CATH Protein Structure Classification database HMMs (CATHDBs) that were specific for importer families in the Transporter Classification Database and found in BGCs included permeases for sugars, oligopeptides, and iron siderophores (**Fig 2c**). 11% of BGCs with a transporter also contained a substrate binding protein. Among the substrate binding proteins we searched for, the most common contained domain was *Peripla_BP_2*, also found in the *E. coli* B12 importer complex BtuCDF, and variants of this protein are specific for siderophores and cobalamin (Berntsson et al. 2010). We also observed many substrate binding proteins with specificities predicted to include carbohydrates, oligopeptides, and peptide uptake (Berntsson et al. 2010) (**Fig 2b**).

In the co-occurrence data, we observed pairing of different substrate binding proteins with different transmembrane domains. *Peripla_BP_2* positively correlated strongly with *FecCD* and *TonB_dep_Rec*, genes known to be involved in siderophore uptake. *BPD_trans_1* co-occurred with either *SBP_bac_5* or *SBP_bac_1*, while *BPD_transp_2* co-occurred with *Peripla_BP_4*. The *ABC2_Membrane_2* domain loosely correlated with *SBP_bac_3*, while the largely uncharacterized *MMPL* transmembrane domain correlated with *SBP_bac_8*. Taken together, these results show a logical organization of importer-specific transporter domains within BGCs that may be involved in either siderophore uptake, precursor uptake, or other roles. Regardless of the substrate specificity of these proteins, care must be taken when assuming that a transporter in a BGC is definitively for export of the matured product.

### Prediction of siderophore and antibacterial activity from biosynthetic transporters

Because transporters are required for the ecological functions of, and are in close contact with, biosynthesized specialized metabolites, we used machine learning to test if transporter classes were predictive of BGC-synthesized metabolite structures and functions. We noticed that metabolite activity labels in MIBiG were strongly associated with phylogeny: 83% of antibiotic BGCs were from gram-positive bacteria while only 40% of siderophore BGCs were from gram-positive bacteria in the dataset of curated MIBiG BGCs. Although this pattern may indicate that the incidence of these functions varies substantially with phylogeny, it is also possible that it reflects a sampling or culturability bias. To reduce the impact of this potential bias, we created separate training and testing datasets for activity prediction for gram-positive and gram-negative organisms.

Using our curated set of BGCs with transporters from the MIBiG 2.0 database, we tested for associations between BGC transport genes and metabolite functions. We generated two activity classification tasks: (a) distinguishing siderophores (including known ionophores) (n=16 gram-positive, n=24 gram-negative), from non-siderophores (n=142 gram-positive, n=42 gram-negative), and (b) distinguishing antibiotics and antifungals (n=131 gram-positive, n=27 gram-negative) from non-antibiotics (n=57 gram-positive, n=37 gram-negative). We observed several statistically significant (Fisher’s exact test; *q*<0.05) associations in the distribution of transporter types between both the siderophore and other activity classes. Among gram-positive bacteria, 60% of siderophore BCGs contained the SBP *Peripla_BP_2* and 55% contained the *FecCD* importer, while no BGCs with other activities had either (**Fig 3a; Fig S1**). The situation was similar for gram-negative siderophore BGCs. The TonB-dependent receptor (completely absent from gram-positive bacteria) was the strongest signal, found in almost 80% of gram-negative siderophore BGCs with a transporter and never in BGCs with other activities.

**Figure 3:**
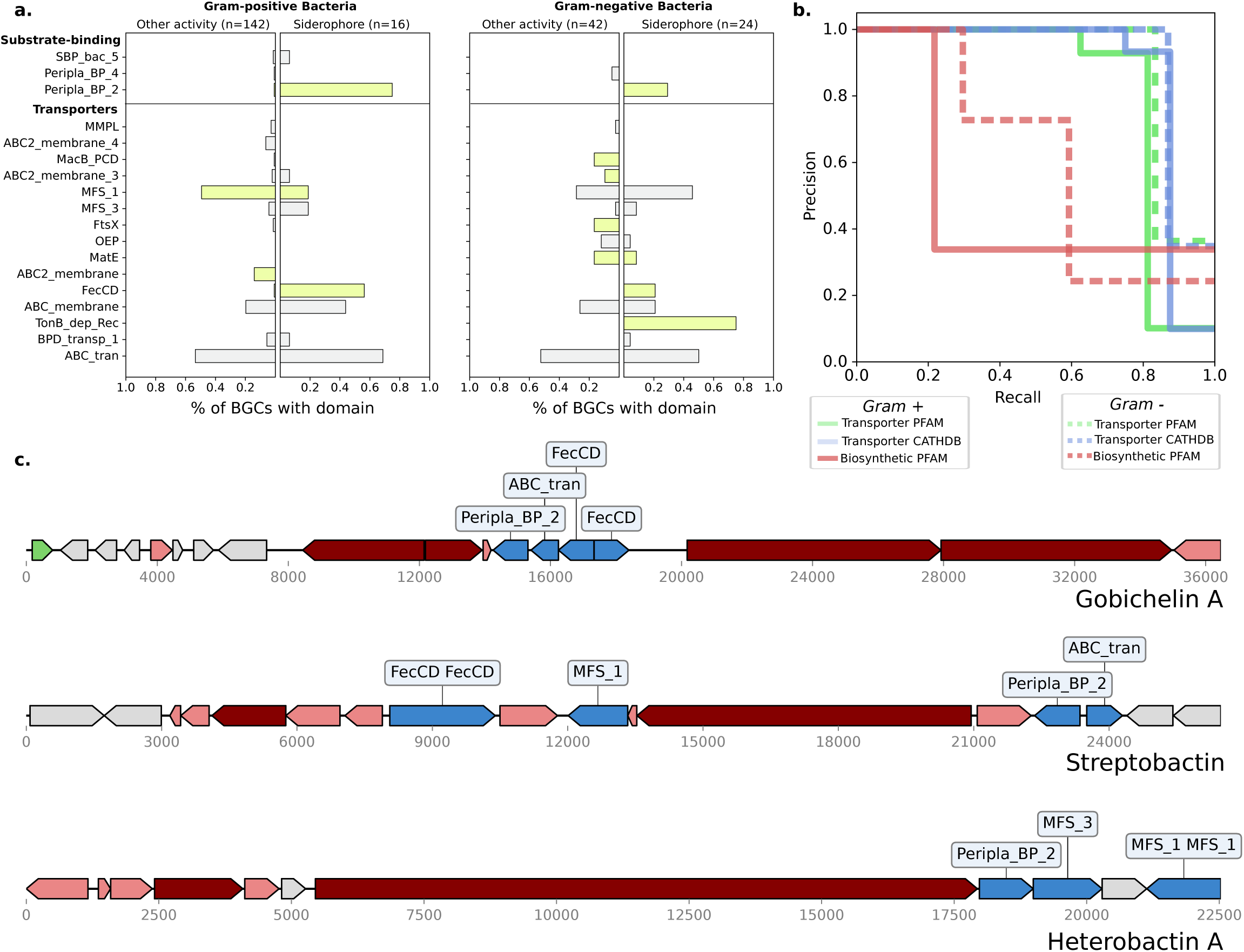
Transporter domains are predictive of siderophore BGCs. (a) The frequencies on common transporter PFAM domains across siderophore BGCs and BGCs of other known activities in gram-positive and gram-negative bacteria. Bars in green were significantly different in frequency between the two classes (Fisher’s exact test; q<0.05) (b) Precision-Recall curves for 2-layer decision trees classifying siderophore BGCs using PFAM transporter, CATHDB transporter, and PFAM biosynthetic gene features in gram negative and gram-positive bacteria. (c) Examples of three siderophore BGCs without activity labels in MiBiG 2.0, which could be identified using transporter frequencies. Transporter genes are colored in blue, core biosynthetic genes (NRPS and PKS) are colored in dark red, accessory biosynthetic genes in light red, and regulatory genes in green.

To assess siderophore predictability from BGC gene content, we used simple decision trees with only two layers applied to different feature sets of protein domain annotations-transport-affiliated PFAMs, transport-affiliated CATHDB HMMs, and biosynthetic PFAMs. To avoid issues with class imbalance, we report precision and recall on the siderophore class, as siderophore prediction requires searching for a minority class (siderophores) within a background of mostly non-siderophores. With the transport-only features, we found that just with two gene decisions, it is possible to achieve 100% precision with over 80% recall using either PFAM or CATHDB transporter annotations for gram-negative siderophores, and 100% precision with over 80% recall using CATHDB transporter annotations for gram-positive siderophores (**Fig 3b; Supplementary Table S5**) On the other hand, when using all biosynthetic annotations within BGCs we found that 2-layer decision trees trained on biosynthetic genes performed substantially worse at predicting siderophore activity than those trained on transporter genes (**Fig 3b; Supplementary Table S5**). We further validated our results by training LASSO linearized regression models, which do not model interactions between features. These models obtained a slightly improved area under the precision-recall curve, indicating that very simple transporter patterns are highly predictive of whether a BGC is siderophore producing or not in our dataset (**Fig 3b**). Transporter features predictive of siderophores were consistently selected by LASSO across stratified cross-validation repeats, giving evidence that these patterns are robust (**Fig S2**). The top predictive biosynthetic features of siderophores were the lucA/lucC protein family (used by antiSMASH to label siderophores and which is known to be involved in aerobactin biosynthesis) and condensation domains (likely to capture non-ribosomal peptide siderophores), but predictive effect sizes were smaller than those for transporters (**Fig S2**).

Using our models for siderophores, we attempted to predict siderophore classes for all gene clusters that have no annotated function in MIBiG2 (and that we had not already hand curated). We searched for gene clusters containing the siderophore-predictive genes *FecCD* and *Peripla_BP_2* and found 6 additional BGCs with no annotated activity, three of which were experimentally validated by the literature to be siderophores (Matsuo et al. 2011; Y. Chen et al. 2013; Carran et al. 2001) (**Fig 3c**). The remaining three identified BGCs were false positives. One of them, the BGC for the antibiotic Ficellomycin, only contained these transport genes in flanking regions not known to be involved in biosynthesis (Liu et al. 2017) while the herbimycin A BGC contains the transporters genes in the reverse reading frame from the BGC, separated by an unusual 16 Kbp intergenic region with only 7% coding density, and the genes appear to be fragmented (Rascher et al. 2005). The other false positive was Lividomycin, an antibiotic that does seem to be a rare non-siderophore BGC with *FecCD* and *Peripla_BP_2* transporters in the MIBiG2.0 database.

Classifying antibiotics and antifungals from either transporters or biosynthetic genes proved more challenging than classifying siderophores (**Supplementary Table S5; Fig S3**). In gram-negative bacteria, we observed positive associations between the MacB-FtsX tripartite efflux pump with antibacterial or antifungal activity (Fisher’s exact test; *q*<0.05). MacB and associated components (FtsX and OEP) were positively associated with antibiotic activity by LASSO logistic regression. This result was stable across both cross-validation folds and repeats of cross-validation (**Fig S3**). We found that 7 out of 27 gram-negative antibacterial BGCs contained a MacB, while no BGCs in our classes of other activities contained MacB. Although MacB is involved in export of the siderophore Pyoverdine (for which there is not an accurate BGC in MiBiG) in *Pseudomonas* (Greene et al. 2018a), in general MacB may be a strong indicator of antibacterial activity for a BGC. Previously, we identified a number of MacB-FtsX exporters in BGCs from novel Acidobacteria (Crits-Christoph et al. 2018), possibly indicating a role in antibacterial activity for these BGCs. LASSO effect sizes for individual biosynthetic genes were substantially lower (**Fig S3**).

### Association of biosynthetic transporter classes with the molecular weights and lipophilicity of their putative substrates

We next hypothesized that transporter classes could be predictive of other molecular features beyond functional activity. There was no strong correlation between the molecular size of the metabolite produced and Gram status of the bacteria encoding each corresponding BCG in the MiBiG dataset. Thus, we tested for differences in transporter classes in BGCs producing metabolites that were (a) less than and (b) greater than 1000 Da in size across all bacteria. There was a striking difference in the frequencies of some transporters between BGCs with different metabolite molecular weights (**Fig 4a**). The strongest difference was in the distribution of MFS transporters - found in 57% of BGCs with products under 1000 Da, but only 14% of those over 1000 Da (Fisher’s exact test; *q*<0.05). The 95th percentile of metabolite molecular weights for clusters with *MFS_1* and without *ABC_tran* was 1082 Da, which may be approaching a biological limit for the molecular weights of substrates for these transporters. Conversely, the ATP-dependent *ABC_tran* domain was found in 89% of BGCs producing high molecular weight compounds but only 42% of those producing low molecular weight compounds (Fisher’s exact test; *q*<0.05). *MacB/FtsX* and the rarer transmembrane domains were also associated with higher molecular weight compounds. In particular, *ABC2_membrane_4* was most associated with high molecular weight compounds, with almost all of the BGCs in which it is found in producing compounds over 1500 Da in size (**Fig 4c; Fig S4**). After training both LASSO logistic regression and 2-layer decision tree models to classify whether produced molecules are greater than 1000 Da, we found that transporter genes were able to distinguish large from small metabolites with moderate precision and recall (**Supplementary Table S5**) and an AUPRC of up to 42%, while biosynthetic genes performed similarly (**Fig 4b**). Top transporter features consistently had larger effect sizes than top biosynthetic features, indicating that transporter-based features provided clearer signals. This result was stable across both cross-validation folds and repeats. (**Fig S4**).

**Figure 4:**
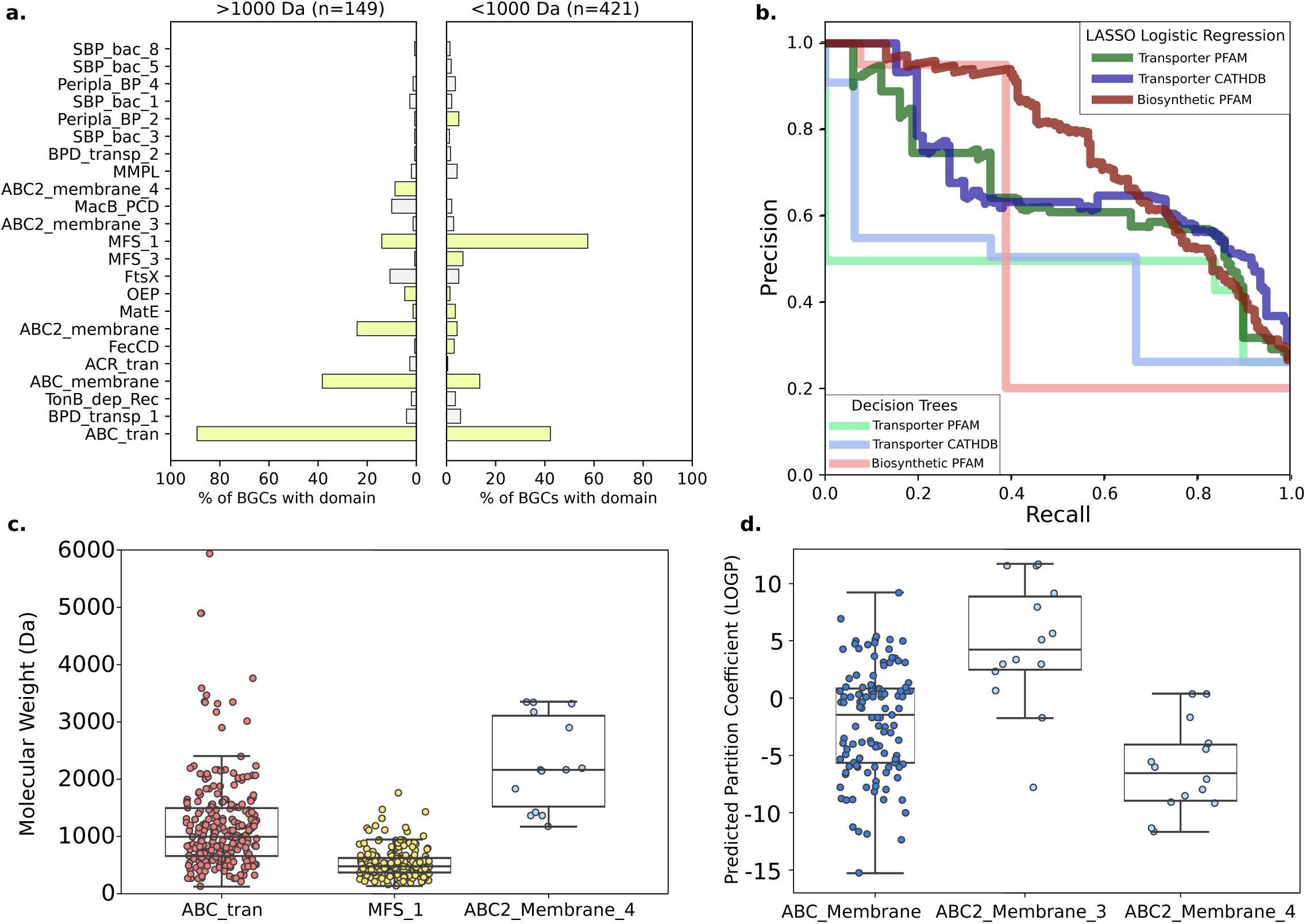
Transporter domains associated with molecular size and partition coefficient. (a) The frequencies of common transporter PFAM domains in BGCs that synthesize metabolites larger than 1000 Da (left) and smaller than 1000 Da (right). Bars in green were significantly different in frequency between the two classes (Fisher’s exact test; q<0.05) (b) Precision-Recall curves for 2-layer decision trees and LASSO logistic regression models classifying BGCs producing metabolites >1000 Da using PFAM transporter, CATHDB transporter, and PFAM biosynthetic gene features. (c) The distribution of metabolite molecular weights synthesized by BGCs with at least one NBD-binding ABC transporter domain, at least one MFS domain, and the ABC2_Membrane_3 transmembrane domain. (d) Predicted partition coefficients (log P) for metabolites synthesized by BGCs that contain at least one variant of three different ABC transporter transmembrane domains.

The biosynthetic protein family most associated with high molecular weight metabolites was *Glycos_transf_2*, likely due to the addition of sugar groups to metabolites by these enzymes in BGCs.

It has previously been reported that transporters can be specific for compounds with similar hydrophilicity (Rempel et al. 2020). The lipophilicity of a metabolite is often considered critical for its success in clinical development for human therapeutics (Arnott and Planey 2012). With LASSO logistic regression, we predicted partition coefficients (log *P*), a measure of lipophilicity, for all of the metabolites in MiBIG, and tested how well metabolite partition coefficients could be predicted by gene content. We found that the presence of 5 transporter classes was significantly associated with increased lipophilicity (Fisher’s exact test; *q*<0.05). In particular, we observed an association between varying ATP-dependent transmembrane domains and log *P*, with *ABC2_Membrane_3* domain co-occurring with BGC-metabolites with a high log *P* (median 4.4) and *ABC2_Membrane_4* domain co-occurring with BGC-metabolites with a low log *P* (median −6.2) (**Fig 4d**). LASSO logistic regression distinguished log P > 0 from log P < 0 with 77% AUPRC (**Supplementary Table S5; Fig S5**). Interestingly, on this task biosynthetic genes were distinctly superior over transporters at prediction of log P, obtaining a 83% AUPRC with a LASSO logistic regression trained on biosynthetic genes (**Supplementary Table S5**).

### Identifying novel siderophore-like biosynthetic gene clusters in the human microbiome

To demonstrate the predictive utility of BGC-associated transporters, we used antiSMASH to annotate all BGCs in sets of genomes assembled from metagenomes from (a) the infant gut microbiome in a study of premature infants from a neonatal intensive care unit (Olm et al. 2019) and (b) a cross-study collation of genomes assembled from multiple human gut studies (Nayfach et al. 2019). We searched these BGCs for the two sets of transporter classes that can achieve near 100% siderophore specificity in the MiBiG database - (a) *Peripla_BP_2* and *FecCD* in gram-positive bacteria, and (b) *Peripla_BP_2, FecCD*, and *TonB_dep_Rec* in gram-negative bacteria. We annotated BGCs with either all of the first two genes or all of the latter three genes as “siderophore-like”, i.e., BGCs that putatively produce siderophores. We identified 1442 BGCs with siderophore-like transporter classes (**Fig 5a; Supplementary Table S6**) and then grouped them into novel gene cluster families using BiG-SCAPE, resulting in 75 siderophore-like gene cluster families (**Fig 5b**). 23% of microbial genomes from the neonatal infant gut microbiomes had siderophore-like BGCs, but only 3% of those assembled from adult gut microbiomes did. Siderophore-like BGCs were identified across a range of bacterial genera, but the vast majority of which were from the *Staphylococcus* or *Enterobacteriales*, which are known to be in high abundance in the neonatal gut microbiome. The genera with the most siderophore-like BGCs were *Staphylococcus* and *Klebsiella*, common hospital-acquired pathogens of neonates. Five of the 75 identified siderophore-like BGC families included a known representative gene cluster in MiBIG, and four of the known representatives were siderophores - again pointing to the specificity of these transporter classes.

**Figure 5:**
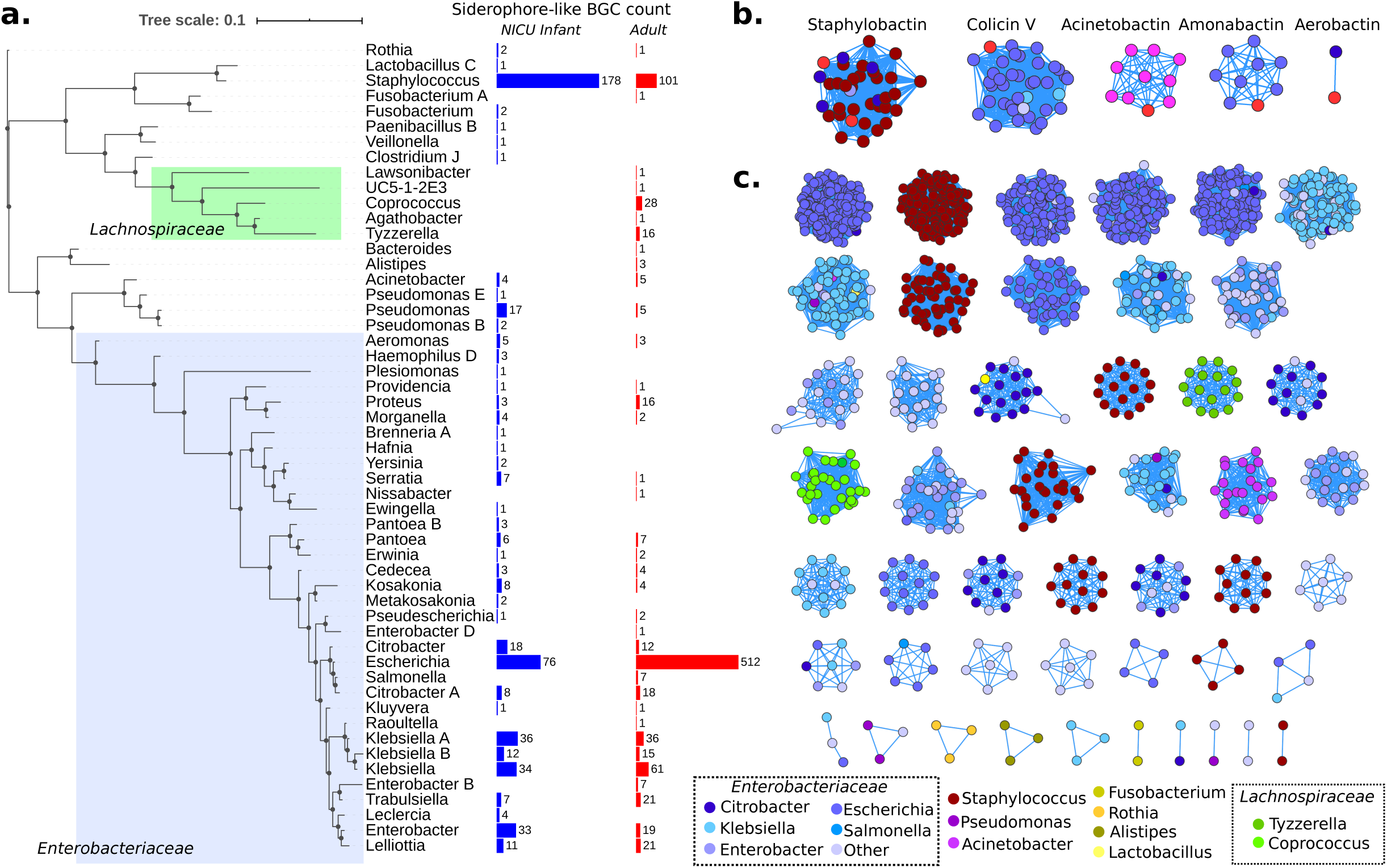
BGCs with siderophore-like transporters from human gut microbiomes. (a) Concatenated ribosomal protein tree (collapsed to the genus level) for high-quality genomes from the infant and adult gut microbiomes that encode siderophore-like BGCs. On the right are counts of siderophore BGCs from infant gut genomes (blue) and adult gut genomes (red). (b) Gene Cluster Families of BGCs containing known siderophores (bright red) and human-microbiome derived BGCs with siderophore-like transporters. BGCs are connected by similarity to other BGCs in the same gene cluster family, calculated using BiG-SCAPE. (c) Families of siderophore-like BGCs without any similarity to existing known BGCs. BGCs in the network are colored by the taxonomy of the genome of origin.

The rest of the siderophore-like gene cluster families that were identified had no closely characterized representative, indicating that there is likely capacity for production of multiple novel siderophores in the human gut microbiome (**Fig 5c**). In the adult gut microbiome samples, these novel siderophore-like gene clusters were identified in phylogenetically diverse genomes, particularly including members of the *Lachnospiraceae*, often considered to be important commensals in the human gut (Duvallet et al. 2017; L. Chen et al. 2017). Most siderophore-like BGCs were in large gene cluster families with other BGCs which also contained the same set of transporter hits.

## Discussion

We uncovered several strong associations between transporters within characterized BGCs and features of the corresponding BGC-synthesized metabolites. With regards to prediction of metabolite activity, we quantified the specificity of TonB-dependent receptors, *FecCD*, and Periplasmic-binding protein 2 for siderophore-producing BGCs. This complements existing literature indicating that genes in these families are specific for siderophore import in both gram-positive and gram-negative bacteria (Chu et al. 2010). We also identified a putative association between the MacB tripartite efflux pump and antibacterial/antifungal activity. Based on these findings, a strategy of targeting novel BGCs containing MacB for characterization may be useful for antibiotic prospecting. In addition to activity prediction, we used metabolite structural information in MiBiG in order to predict metabolite molecular weight and lipophilicity from BGC gene content. We discovered a strong relationship between the transporters in characterized BGCs and the molecular weight of their synthesized metabolites. The strong dichotomy between ATP-dependent transporters (utilizing the *ABC_tran* nucleotide binding domain) and *MFS* family transporters points towards required ATP-dependence for transporting metabolites larger than 1000 Da. We also identified relationships between two understudied membrane components (*ABC2_Membrane_3* and *ABC2_Membrane_4*) and substrate log *P*, possibly indicating tradeoff in membrane domains for molecules of different chemical properties. Future phylogeny-based subdivision of these families may improve upon general protein family annotations to increase the predictive power of transporter substrate characteristics.

There are multiple caveats to our work. Molecular activity of specialized metabolites based on functions proven in the laboratory may be very different from the ecological roles that metabolites play in natural settings (van der Meij et al. 2017; Kramer, Özkaya, and Kümmerli 2020; Behnsen and Raffatellu 2016). Further, many BGCs may produce multiple variants of a metabolite (Fischbach and Clardy 2007), only some of which may be reported. There may also be reference-database biases in our gene searches - while they are sensitive, it is possible that phylogenetically divergent microbes use transporter genes that are not hit by our sequence models. Finally, as reported, a significant proportion of BGCs contain no transporter at all or a transporter gene genomically adjacent to a BGC may not be functionally linked. Despite these caveats, it appears as though transporter genes provide simple and strong signals for inferring both activity and chemical properties of metabolites produced by BGCs.

Siderophores are both considered critical pathogenicity factors for many human-associated microbes (Weakland et al. 2020) and are also known to facilitate interactions with other microbes and the innate immune system in the human gut microbiome (Behnsen and Raffatellu 2016; Holden et al. 2016; Zhu et al. 2020; Lam et al. 2018). Therefore, being able to annotate genes for the production of siderophores across diverse bacterial species may be critical for understanding the distribution of virulence factors, yet it is difficult to do using traditional annotation pipelines alone. We observed a high prevalence of siderophore-like BGCs in bacterial genomes from NICU premature infant guts, suggesting that the premature infant gut could be more prone to invasion by pathogens with siderophore virulence factors. Potentially novel siderophore-like BGCs were most consistently found to be encoded in the genomes of members of the *Enterobacteriaceae* and *Staphylococcus* in the premature infant microbiome. Only in the adult microbiome datasets did we identify siderophore-like BGCs in the *Lachnospiraceae*, that are often considered important commensals, indicating that commensal siderophore production may be more common in developed gut microbiomes. Importantly, we identified siderophore-like BGCs in these taxa that are not homologous to known siderophore clusters, indicating that there is still substantial unknown chemical diversity of siderophores, even within well-studied lineages.

In general, here we demonstrated that consideration of transporter genes can aid holistic functional prediction of BGC products. A transporter guided approach could be especially useful for identification of siderophore targets for medical (Nagoba and Vedpathak 2011) and biotechnological applications(Ahmed and Holmström 2014). Given the large diversity of BGCs and that chemical characterization of their products can be time and resource intensive, better functional prediction of BGCs for targeted study can improve selection of targets for antimicrobial discovery and downstream activity tests.

## Methods

### Curation and selection of BGCs and transporter annotations

We parsed the MIBiG 2.0 database of biosynthetic gene clusters metadata and extracted information including host genus, compound count, chemical structures of the metabolite product, known metabolite activities, and the number of open reading frames for each BGC. Using Entrez and NCBI, we assigned the expected Gram status for each BGC based on phylum, coded 0=Gram-negative, 1=Gram-positive, 2=Fungal, 3=other. For the purpose of this manuscript, we only analyzed BGCs from Gram-positive and Gram-negative bacteria. We noticed that the activity labels in MIBiG 2.0 were often incomplete, and manually added a set of antibacterial, siderophore, and antifungal labels derived from the literature (**Supplementary Table S1**). We found that 28 BGCs (1.8%) in MIBiG were unusually large in length, and comparisons to published papers on these BGCs showed that their MIBiG counterparts were overextended in comparison to the validated BGC. For this reason, we eliminated the 28 BGCs over 60 ORFs in length. Using Python and RDKit, we calculated molecular weights and partition coefficients (log *P*) using the algorithm described in (Wildman and Crippen 1999) for all 1,042 MIBiG BGCs with a single associated compound structure. 238 BGCs have more than 1 associated metabolite structure, and these multi-structure BGCs were not used in our structural association analyses. We annotated biosynthetic genes in MIBiG with HMMER *hmmsearch (Eddy 1998)* and *cath-resolve-hits (Lewis, Sillitoe, and Lees 2019)* on a set of the 99 most commonly represented biosynthetic PFAMs in antiSMASH BGCs obtained from (Cimermancic et al. 2014), to generate a counts table of biosynthetic protein families for each BGC (**Supplementary Table S4**).

To obtain a comprehensive overview of the distribution of transport-associated protein domains in biosynthetic gene clusters, we generated two separate feature tables: (a) using CATHDB (Sillitoe et al. 2019) HMMs and (b) using PFAM (“PFAM Database,” n.d.) HMMs. For the first, we downloaded all proteins in the Transporter Classification Database (TCDB) (Saier et al. 2016) and annotated them with *hmmsearch* and *cath-resolve-hits*, using all CATHDB Functional Family HMMs. We then selected all CATHDB HMMs which were represented at least 5 times. We then manually curated this list down to 180 final CATHDB HMMs that were transport-specific. We then calculated the specificities of each CATHDB HMM for TCDB families; 80 were specific for exactly one TCDB family.

For the second set of features, we took all protein sequences with an annotation including “Transport” in the antiSMASH database v2 (Blin, Pascal Andreu, et al. 2019) and annotated these proteins with the Pfam-A set of HMMs using *hmmsearch* and the option --cut_ga. We then selected highly represented HMMs and manually curated this list to be transporter-specific and representative of the major transporter classes in TCDB. We then also selected the PFAM Substrate Binding Protein and Periplasmic Binding Protein HMMs that were represented more than 5 times in MIBiG. When comparing both the PFAM and CATHDB set of HMMs, we found substantial overlap, but the CATHDB set is composed of 166 domain features while the PFAM set only contains 18.

### Machine Learning to predict metabolite structural and functional characteristics

To identify associations between metabolite functional classes and structural properties with BGC gene content, we used traditional statistical tests and different machine learning models. Our classification tasks were (a) siderophore (n=16 gram-positive; n=24 gram-negative) vs other activity (n=142 gram-positive; n=42 gram-negative), (b) antibiotic and antifungal (n=131 gram-positive; n=27 gram-negative) vs other activity (n=57 gram-positive; n=37 gram-negative), (c) metabolite molecular weight > 1000 Daltons (n=149) vs metabolite molecular weight < 1000 Daltons (n=421), and (d) predicted partition coefficient log *P* <0 (n=220) vs log *P* >= 0 (n=350). For the functional classification tasks, we noticed a strong class imbalance with gram-status, so we performed functional classification separately for BGCs from gram-positive and gram-negative bacteria. We first tested for univariate differences in proportional representation of each transporter BGC between classes for classification tests using Fisher’s Exact Test in Python (q<0.05, Benjamini-Hochberg (Hochberg and Benjamini 1990) correction).

We then assessed the predictive power of the three sets of features for each BGC: transporter PFAM HMMs, transporter CATHDB HMMs, and biosynthetic PFAM HMMs. Features were counts of protein families, which were standardized using the StandardScaler function in the scikit-learn package. Given the nature of our study, we used simple models to ensure reliability and interpretability of our results. We fit two classes of machine learning models (a) LASSO-penalized logistic regression, which fits a linear model with a sparsity penalty on weights, and (b) shallow decision trees (of depth one or two), which can classify based on splitting at most two features (Franklin 2005). All models were trained using the Python package scikit-learn. Due to data size and class imbalance, we fit models using repeated, stratified k-fold cross-validation (“RepeatedStratifiedKFold” in scikit-learn) with five repeats and five folds. On each cross-validation split of our data, we computed the area under the precision-recall curve (AUPRC) to evaluate performance for our class-imbalanced tasks. Thus for the final output of this procedure we reported the mean of accuracy, precision, recall, and AUPRC each generated from five repeats of different random 5-fold partitions of the data. We further use these repeated cross-validation splits to see which features are consistently used for classification across repeats.

### Annotating metagenomic siderophore-like BGCs from the human microbiome

To assess the distribution of siderophore-like BGCs in the human microbiome, we downloaded two sets of genomes assembled from metagenomes obtained from the human gut microbiome: 2,425 genomes from a neonatal intensive care unit (NICU) premature infant microbiome (Olm et al. 2019) and 24,345 genomes from a diverse set of mostly adult human cohorts (Nayfach et al. 2019). We ran antiSMASH 5.0 on these genomes and then scanned predicted BGCs for at least two of the PFAMs that were found to be specific for siderophore BGCs (*FecCD, Peripla_BP_2*, and *Ton_dep_Rec*). We dereplicated these BGCs and compared them to known MIBiG siderophore BGCs using the software BiG-SCAPE (Navarro-Muñoz et al. 2020) run with default settings. We then considered BGCs with either set of hits and reported their genomic taxonomic distribution based on the closest BLAST hit representatives of genomic ribosomal proteins to taxonomic genera defined by GTDB (Parks et al. 2018) (minimum percent identity of hits >80%).

## Acknowledgements

This research was supported, in part, by the NIH under awards RAI092531A, R01-GM109454, and R35-GM134922. J.F.B. and Y.S.S. are Chan Zuckerberg Biohub Investigators.

## Data Access

All Python code used in this paper, along with the data analyzed, antiSMASH BGCs, and additional data tables, is available at https://github.com/nickbhat/bgc_tran.

## Supplementary Figure Legends

**S1. Heatmap of PFAM transporter domain abundance in all annotated siderophores in our modified version of the MIBiG database**.

**S2. Distributions of and LASSO logistic regression feature importances of transporter and biosynthetic gene features for prediction of siderophore activity**.

**S3. Distributions of and LASSO logistic regression feature importances of transporter and biosynthetic gene features for prediction of antibacterial activity**.

**S4. LASSO logistic regression feature importances of transporter and biosynthetic gene features for prediction of molecular weights >1000 Da**.

**S5. Precision-recall curves and LASSO logistic regression feature importances of transporter and biosynthetic gene features for prediction of log P > 0**.

## Supplementary Table Legends

**S1. Metadata for all MIBiG BGCs included in this study, including additional hand curated labels for data**.

**S2. Abundances of PFAM transporter domains across all MIBiG BGCs**.

**S3. Abundances of CATHDB transporter domains across all MIBiG BGCs**.

**S4. Abundances of PFAM biosynthetic domains across all MIBiG BGCs**.

**S5. Classification metrics (precision, recall, accuracy, and AUPRC) for all classification tasks in this study**.

**S6. Siderophore-like Biosynthetic Gene Clusters identified in the human microbiome by this study**.

