## Supplementary figures and images for "Transporter genes in biosynthetic gene clusters predict metabolite characteristics and siderophore activity"

### FigS2.pdf

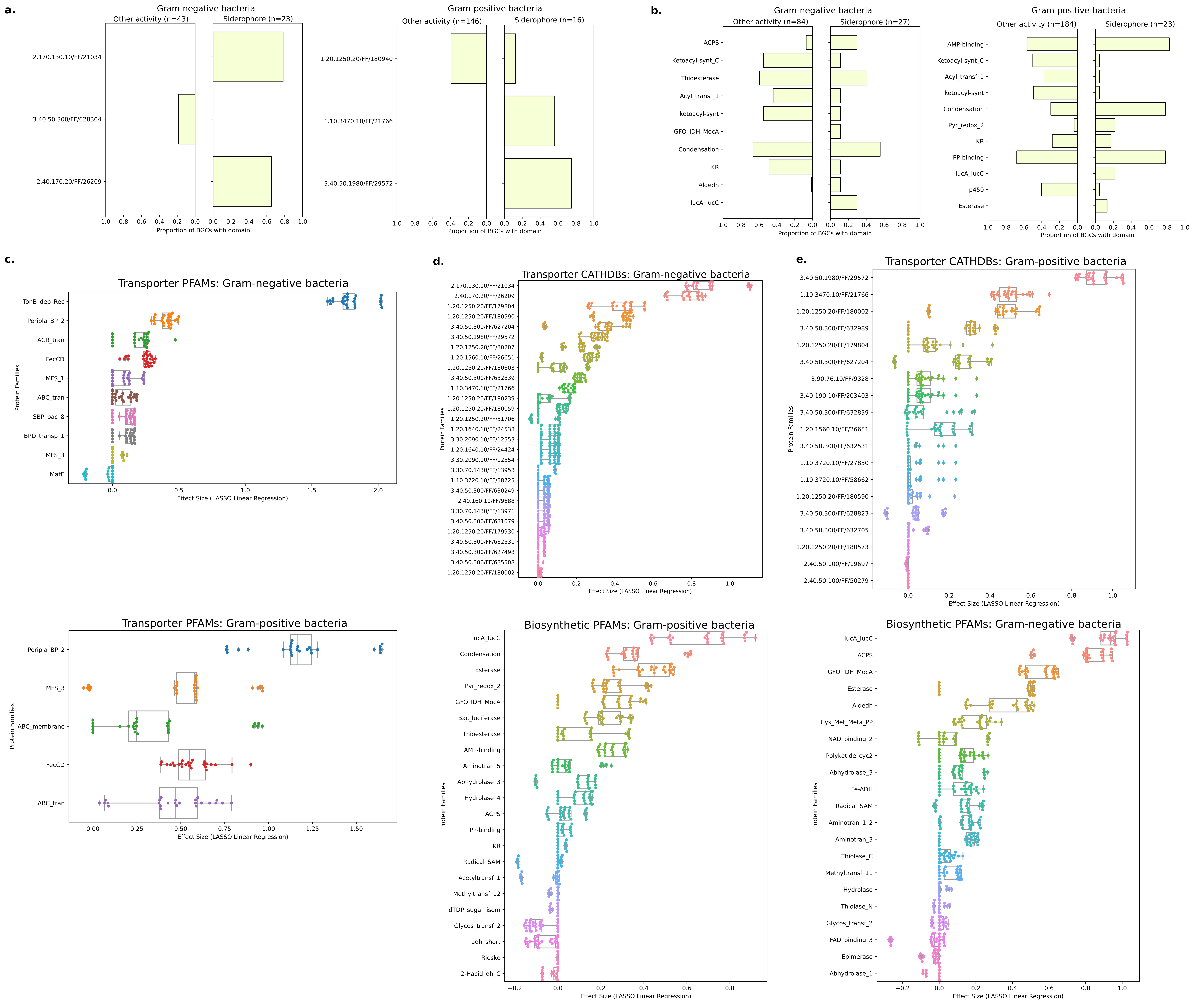

### FigS3.pdf

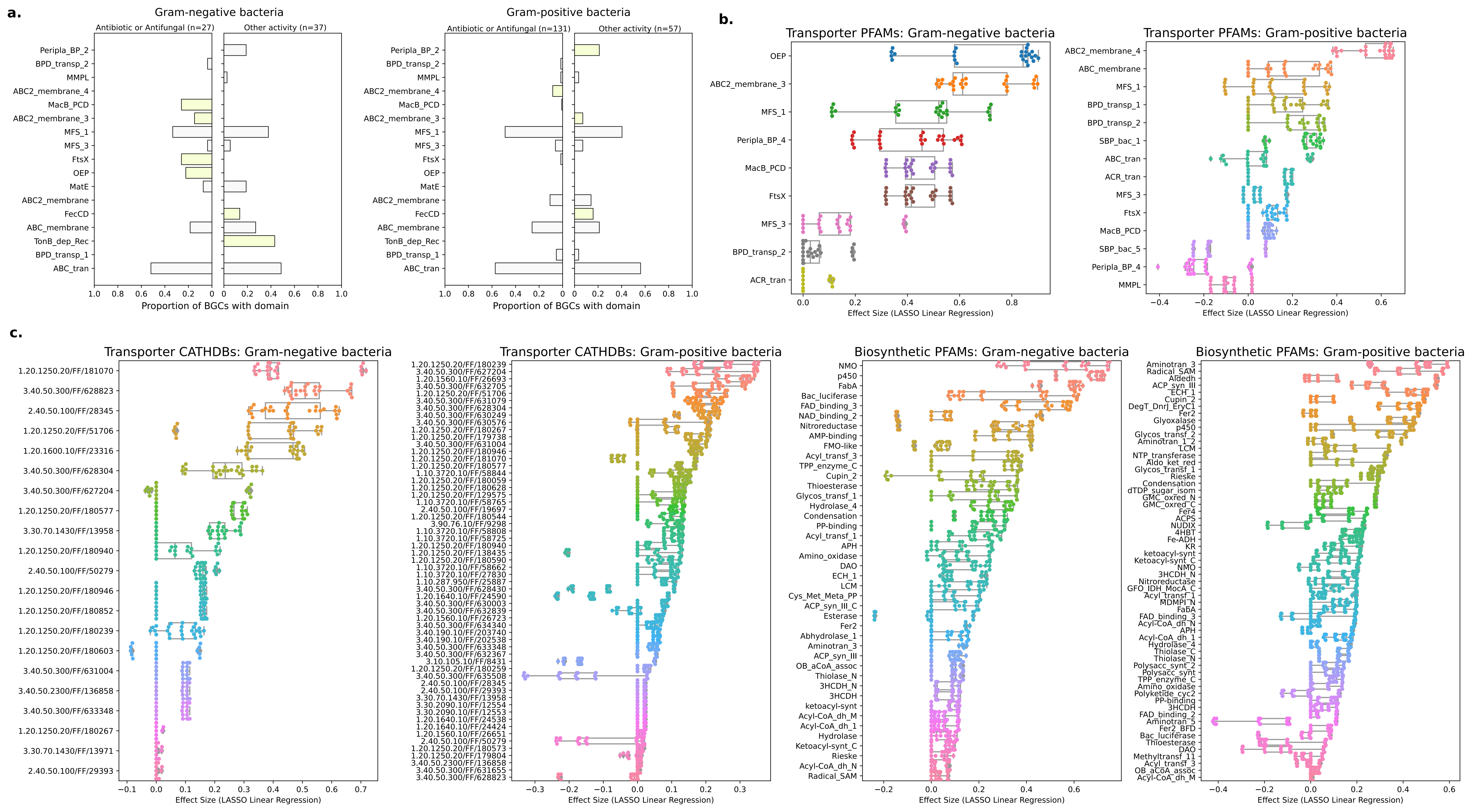

### FigS4.pdf

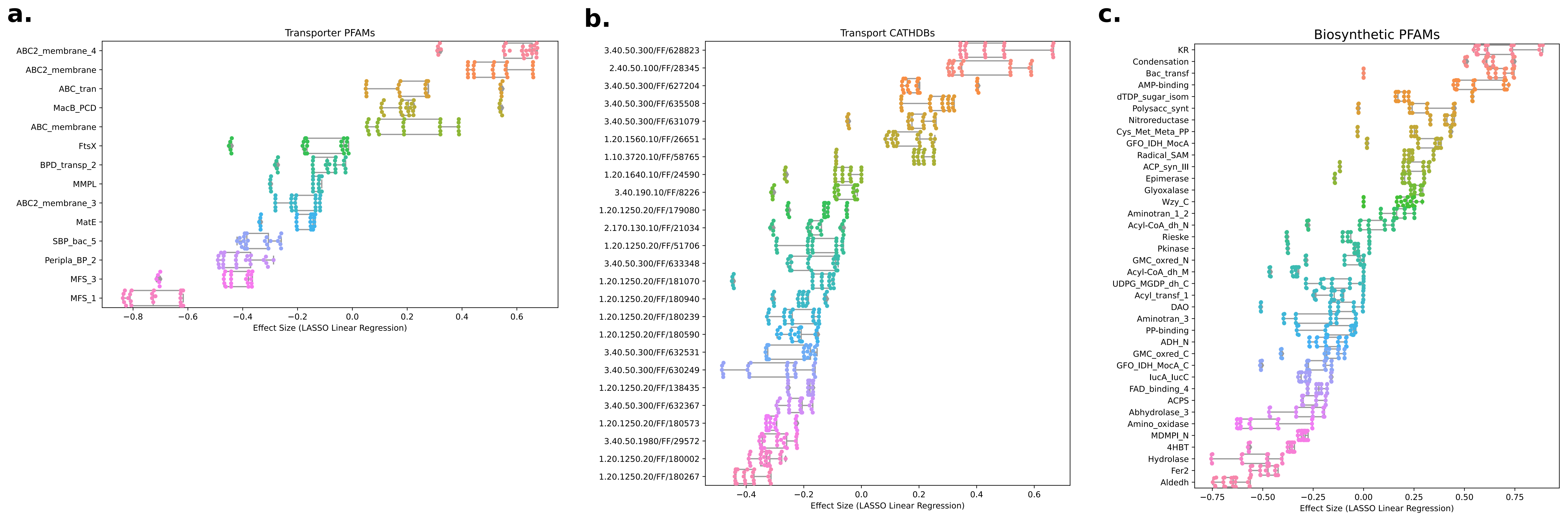

### FigS5.pdf

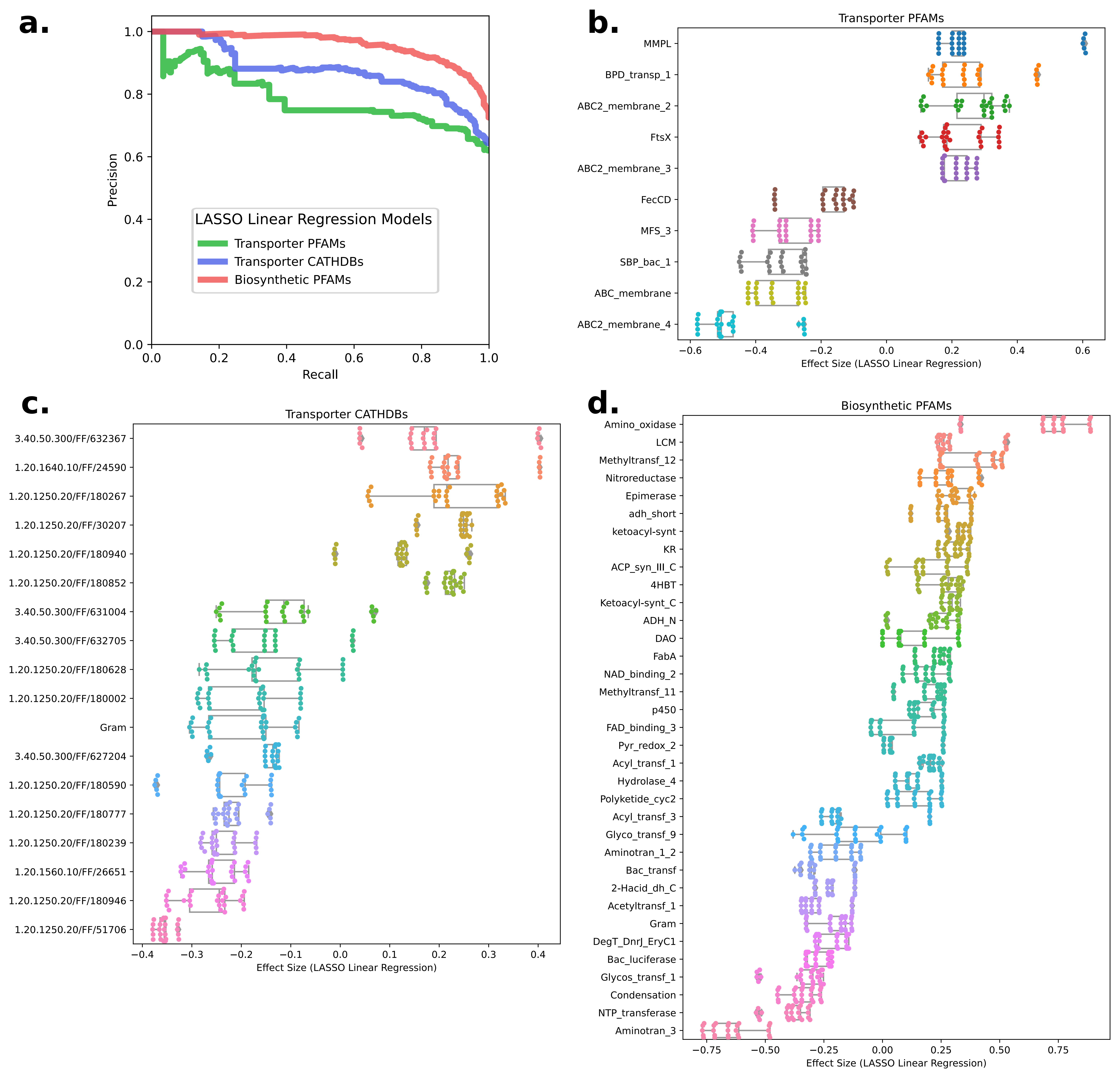

### S1.pdf

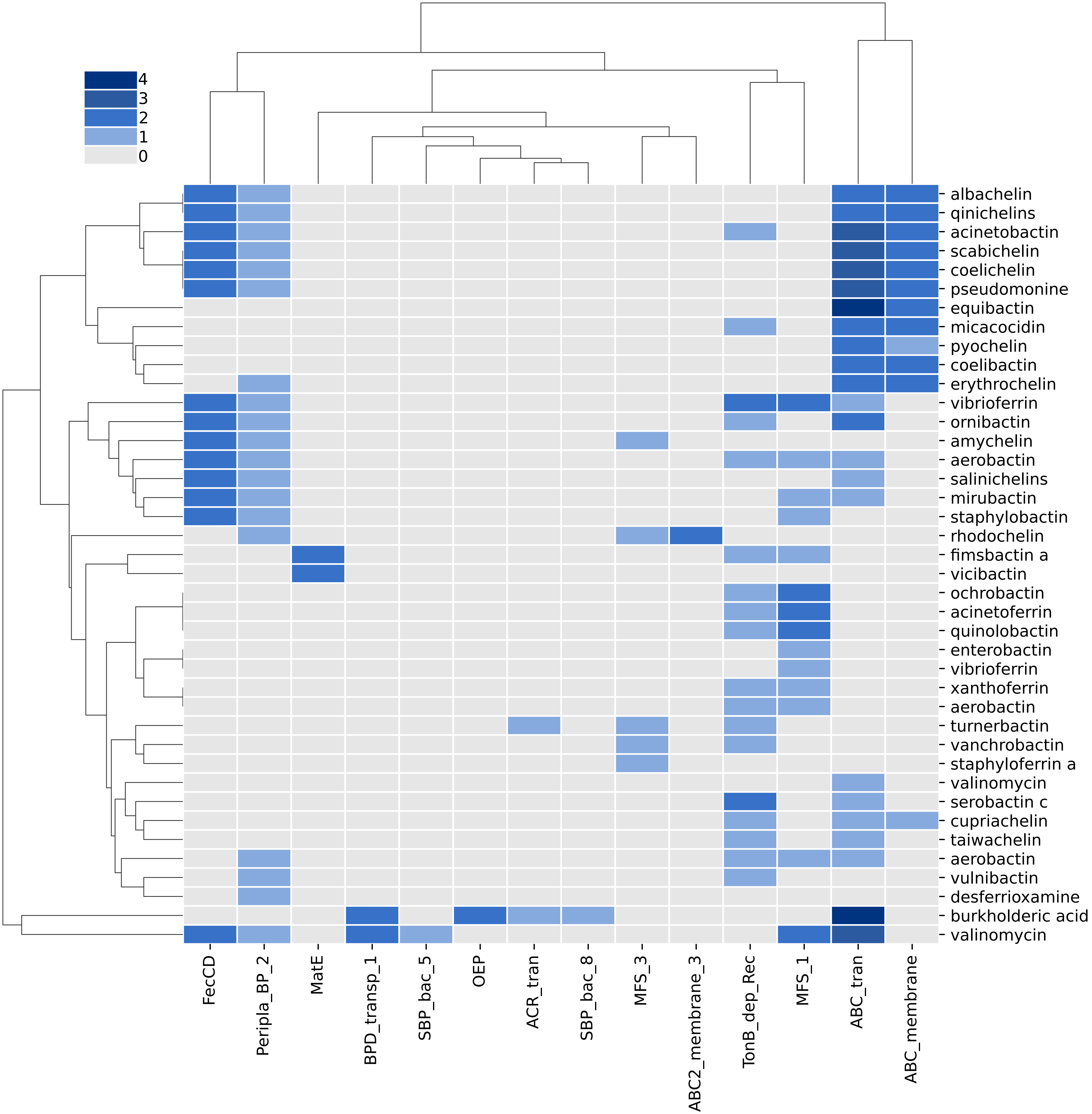
